# Gene discovery for Mendelian conditions via social networking: *de novo* variants in *KDM1A* cause developmental delay and distinctive facial features

**DOI:** 10.1101/028241

**Authors:** Jessica X. Chong, Joon-Ho Yu, Peter Lorentzen, Karen M. Park, Seema M. Jamal, Holly K. Tabor, Anita Rauch, Margarita Sifuentes Saenz, Eugen Boltshauser, Karynne E. Patterson, Deborah A. Nickerson, University of Washington Center for Mendelian Genomics, Michael J. Bamshad

## Abstract

**Purpose:**The pace of Mendelian gene discovery is slowed by the “n-of-1 problem” – the difficulty of establishing causality of a putatively pathogenic variant in a single person or family. Identification of an unrelated person with an overlapping phenotype and suspected pathogenic variant in the same gene can overcome this barrier but is often impeded by lack of a convenient or widely-available way to share data on candidate variants / genes among families, clinicians and researchers.

**Methods:**Social networking among families, clinicians and researchers was used to identify three children with variants of unknown significance in *KDM1A* and similar phenotypes.

**Results:**De novo variants in *KDM1A* underlie a new syndrome characterized by developmental delay and distinctive facial features.

**Conclusion:**Social networking is a potentially powerful strategy to discover genes for rare Mendelian conditions, particularly those with non-specific phenotypic features. To facilitate the efforts of families to share phenotypic and genomic information with each other, clinicians, and researchers, we developed the Repository for Mendelian Genomics Family Portal (RMD-FP). Design and development of a web-based tool, MyGene2, that enables families, clinicians and researchers to search for gene matches based on analysis of phenotype and exome data deposited into the RMD-FP is underway.

## INTRODUCTION

Gene discovery strategies based on exome and whole genome sequencing (ES/WGS) that are agnostic to both known biology and mapping data provide powerful alternatives to conventional approaches to gene identification. Since their introduction in 2010, ES/WGS-based strategies have proven to be disruptive technologies that have rapidly accelerated the pace of discovery of genes underlying Mendelian phenotypes.^1^ For example, the rate of reported gene discovery increased from an average of ~166 per year between 2005 and 2009 to ~236 per year between 2010 and 2014, or an increase of 40% (i.e., ~70 additional reports) per year.^1^ However, this increase in reported discoveries is more modest than we, and perhaps others, anticipated.

Among the myriad factors limiting ES/WGS-based gene discovery, one key challenge is the lack of infrastructure for (1) large-scale standardized phenotypic delineation and comparison of families with Mendelian conditions, and (2) open sharing of sequence data, candidate genes, and putative causal variants between investigators and clinicians. These limitations often result in identification of a putative causal variant or several high-priority candidate variants in an individual who has an unexplained (i.e., causal gene unknown) phenotype, requiring extensive functional experimentation to establish a causal relationship.^2,3^ In clinical settings, this issue frequently manifests as the reporting of a variant of unknown (or uncertain) significance (VUS).^4,5^ In contrast, identification of novel putative causal variants in the same gene in two or more families with the same or similar phenotype strongly supports a causal relationship independent of functional studies.^3^ To this end, sharing phenotypic and genetic information among investigators and clinicians in order to find multiple families with putatively pathogenic variants in the same gene is a straightforward approach to establishing a causal relationship. This is the rationale for developing infrastructure for large-scale release of combined genotypeto-structured phenotype data (e.g. Geno_2_MP^1^), structured phenotype matching^6^ (e.g. PhenomeCentral), gene matching (e.g. GeneMatcher^7^, and variant matching (e.g. GenomeConnect^8^, Decipher/DDD^9,10^), which are being coordinated via efforts such as Matchmaker Exchange^11^.

Few, if any, formal resources for phenotype or gene matching exist that meet the needs of families motivated to identify individuals with similar phenotypes or VUS in the same gene. Instead, families have turned to the internet and social media as a way to share experiences and knowledge with other families and researchers in an effort to more fully leverage the diagnostic potential of clinical genetic testing. Such efforts at internet-driven patient finding (IDPF)^12^ led, for example, to the widely-publicized delineation of a novel disorder of glycosylation caused by loss-of-function variants in *NGLY1*.^13,14^

Inspired by this success,^15^ the parents of a child (Family A) with developmental delay, hypotonia, and multiple minor anomalies, in whom clinical ES^16^ identified *de novo* variants of unknown significance in *lysine (K)-specific demethylase 1A* (MIM 609132; *KDM1A*) and *ankyrin repeat domain-containing protein 11* (MIM 611192; *ANKRD11*), established a website, Twitter account, and Facebook page to publicize these findings. Their goal was to identify other families with similarly affected children and / or VUS in the same gene(s) and recruit researchers to study their child’s condition. Their efforts were successful and within five days led to the identification of another family (Figure 1, Family B; Figure S1) who had a child with similar clinical characteristics and a *de novo* VUS in *KDM1A* (Figure 2). Family A also contacted via e-mail various research groups in the United States investigating the genetic basis of developmental delay. A member of one of these groups made Family A aware of a publication in which a *de novo* VUS in *KDM1A* had been reported in a child with severe non-syndromic intellectual disability and unaffected parents (Family C)^17^. Literature searches by the diagnostic laboratory and clinicians for Families A and B, and Family A themselves had failed to identify Family C in part because the information on VUS, including the one in *KDM1A*, identified in the cohort was listed only in the supplementary materials of the manuscript.

**Figure 1.**
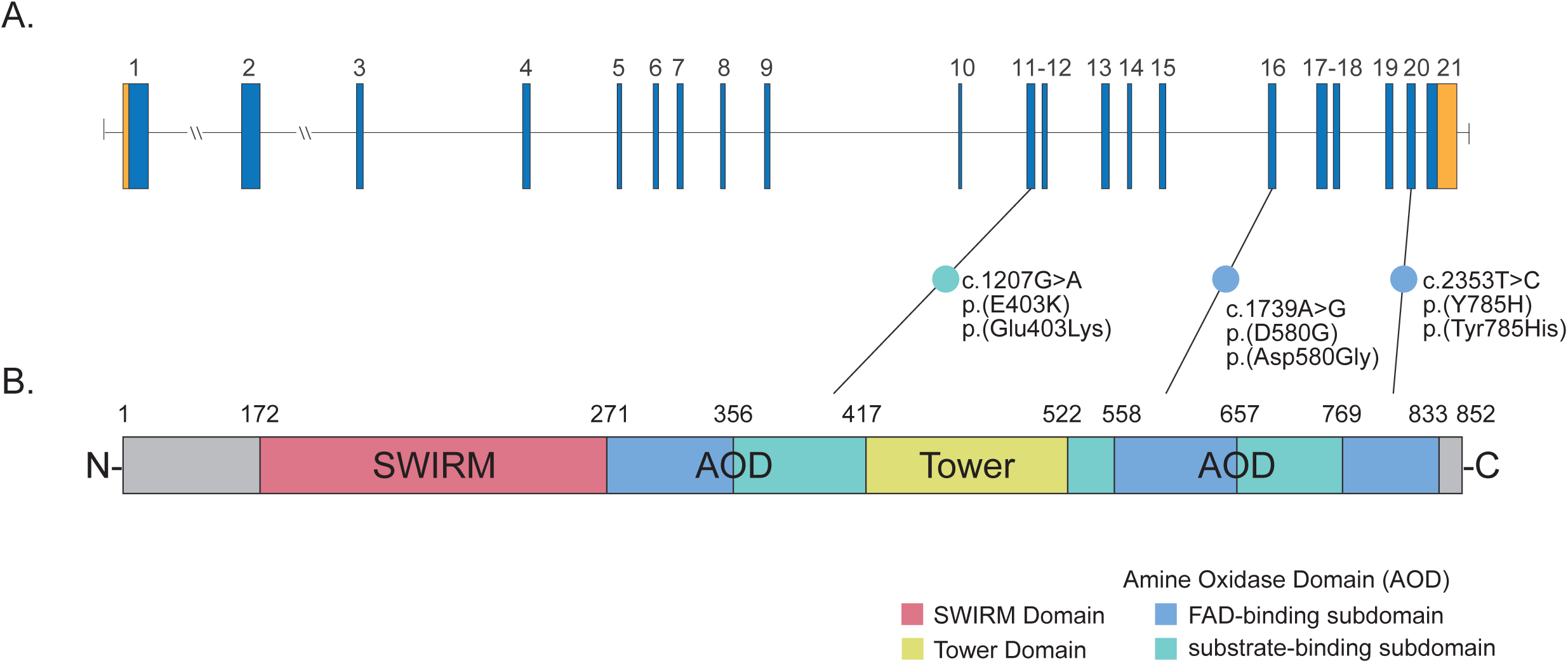
Phenotypic characteristics of children with a mutation in *KDM1A*. *Note: This figure has been omitted from the submission of this manuscript to bioRxiv.org at the request of their board.* All three individuals (A-C) with a mutation in KDM1A share a prominent forehead, slightly arched eyebrows, elongated palpebral fissures, a wide nasal bridge, thin lips, and wide-spaced teeth. Case identifiers correspond to those in Table 1, where a detailed description of the phenotype of each person is provided. C-1 and C-2 are pictures of the same child at 3 years 8 months and 8 years of age, respectively.

**Figure 2.**
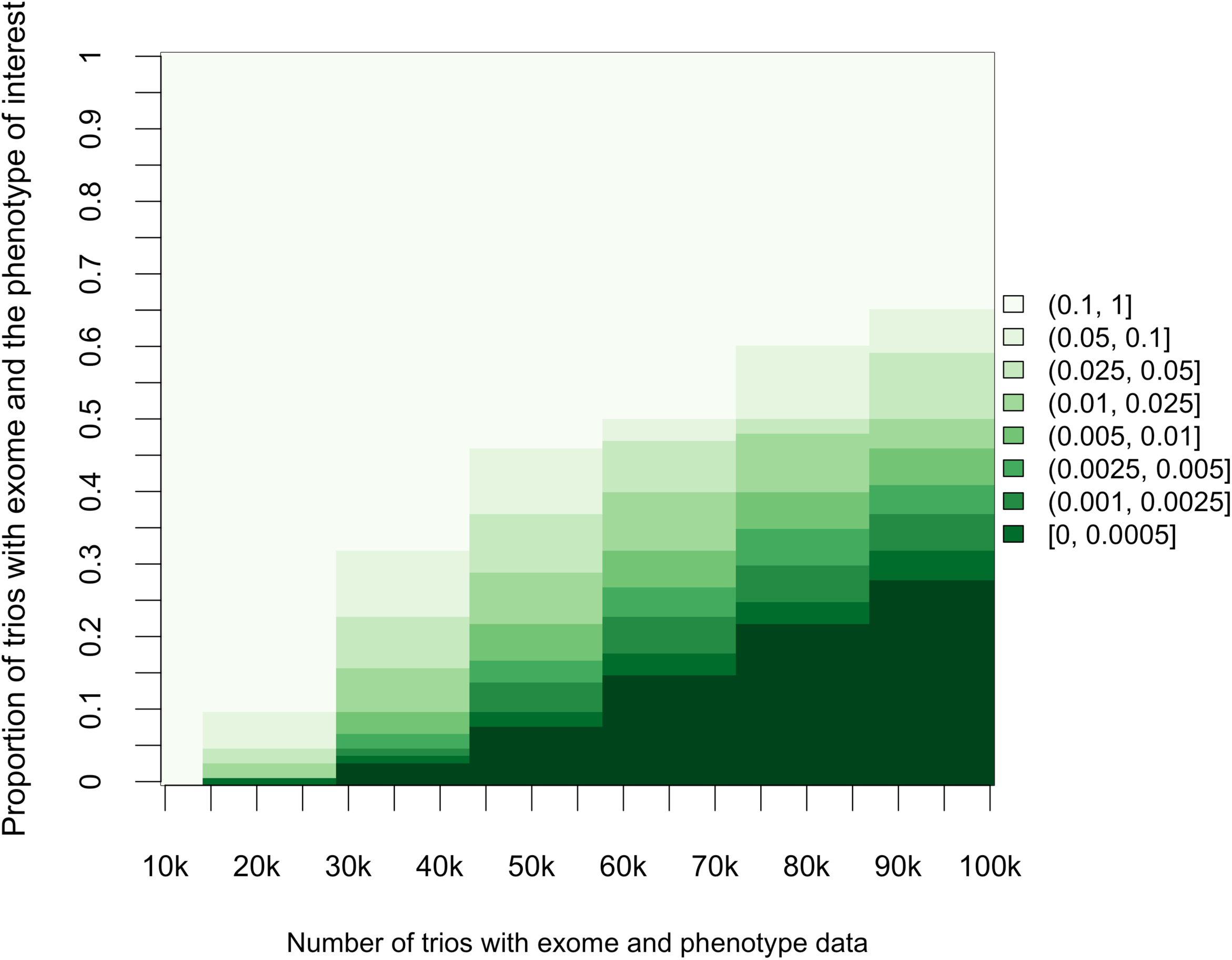
Genomic structure of *KDM1A*, predicted KDM1A protein, and spectrum of mutations that cause developmental delay. A) *KDM1A* is composed of 21 exons including protein-coding (blue) exons and non-coding (orange) exons. Lines with attached dots indicate the approximate locations of the three different *de novo* variants that we report to underlie developmental delay. The color of each dot reflects the domain/subdomain containing the corresponding mutated residue. B) Protein domain structure of KDM1A. KDM1A has three domains—SWIRM (pink), Amine-Oxidase Domain (AOD, blue and teal), and Tower (yellow)—as well as an unstructured N-terminal flexible region and C-terminal tail (gray). The amine-oxidase domain (AOD) is comprised of two subdomains, the FAD-binding and substrate-binding functional subdomains. The active site cavity of KDM1A is within the substrate-binding subdomain and is required for KDM1A to demethylate H3K4me1/2 and repress transcription. Both the Tower and SWIRM domains have been shown to be necessary for the catalysis of histone demethylation by KDM1A.

Subsequently, the parents of Family A (P.L. and K.M.P.) sought out the expertise of investigators at the University of Washington Center for Mendelian Genomics (UW-CMG) to help delineate the condition, confirm that variants in *KDM1A* were likely to be causal, and report the findings to the human genetics community at large. This experience with gene discovery for a Mendelian condition via social networking prompted design and preliminary development of a web-based portal (MyGene2) that will be accessible via the UW-CMG, through which families can submit phenotypic information and sequence data (e.g., VCF and BAM files) to be warehoused and made accessible to researchers worldwide in order to facilitate more universal IDPF.

## MATERIALS AND METHODS

Studies were approved by the University of Washington and the University of Zurich Institutional Review Boards and consent to publish photographs was obtained. For two of the three families (Figure 1, Table 1, Figure S1; Families A and B), clinical ES was performed at GeneDx (Gaithersburg, MD) using the Agilent SureSelect XT2 All Exon V4 target. Both families requested BAM files from GeneDx and upon receipt, each family transferred the files to the UWCMG where they were reprocessed using a standard pipeline as previously described^18^. Reads were aligned to a human reference (hg19) using the Burrows-Wheeler Aligner (BWA) 0.6.2. All aligned read data were subjected to: (1) removal of duplicate reads (Picard MarkDuplicates v1.70) (2) indel realignment (GATK IndelRealigner v1.6-11-g3b2fab9); and (3) base quality recalibration (GATK TableRecalibration v1.6-11-g3b2fab9). Variant detection and genotyping were performed using GATK UnifiedGenotyper (v1.6-11-g3b2fab9). Variant data for each sample were flagged using the filtration walker (GATK) to mark sites that were of lower quality and potential false positives (e.g. strand bias≥-0.1, quality scores (QUAL≤50), allelic imbalance (ABHet>0.75), long homopolymer runs (HRun>4), and/or low quality by depth (QD<5).

**Table 1.**
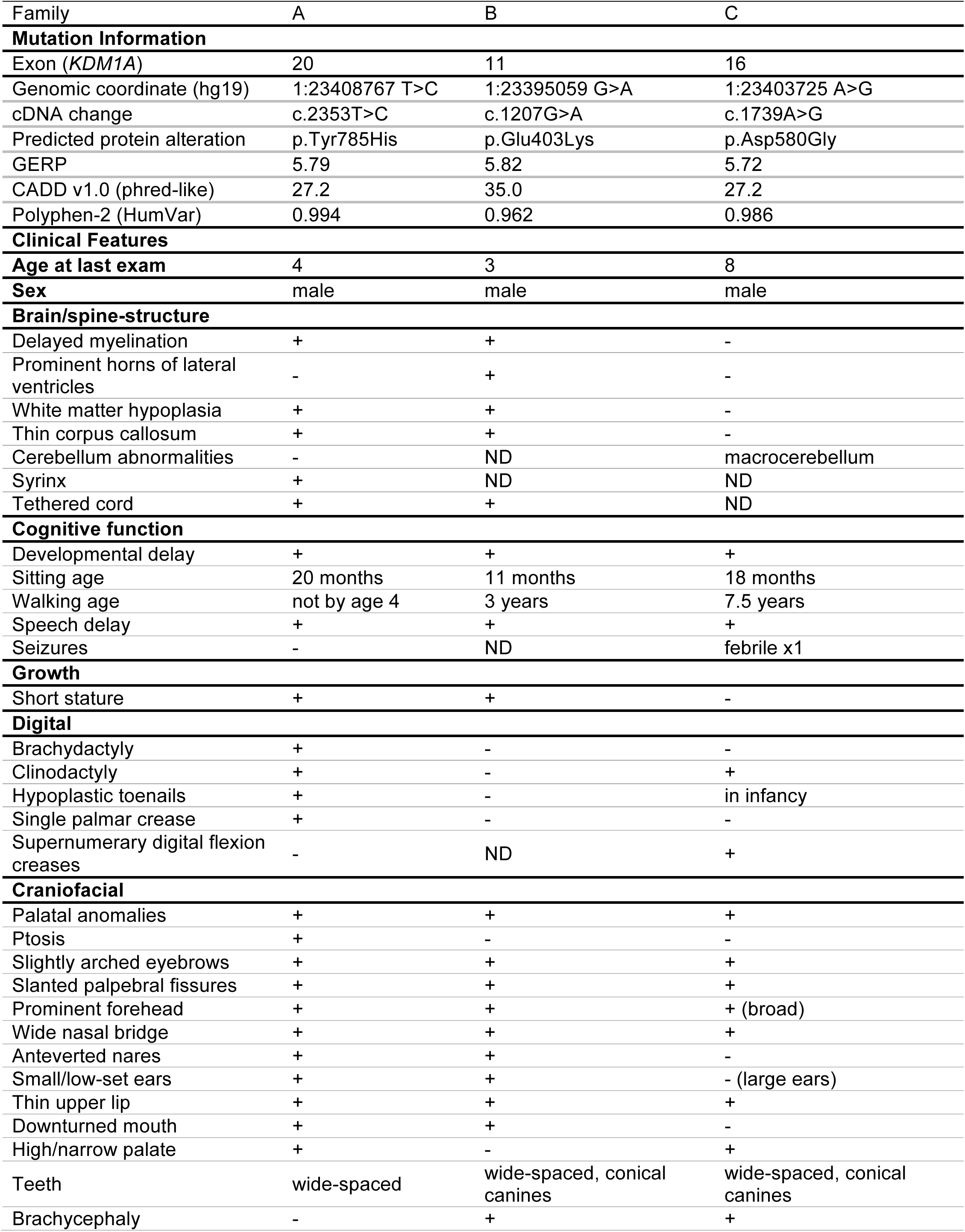
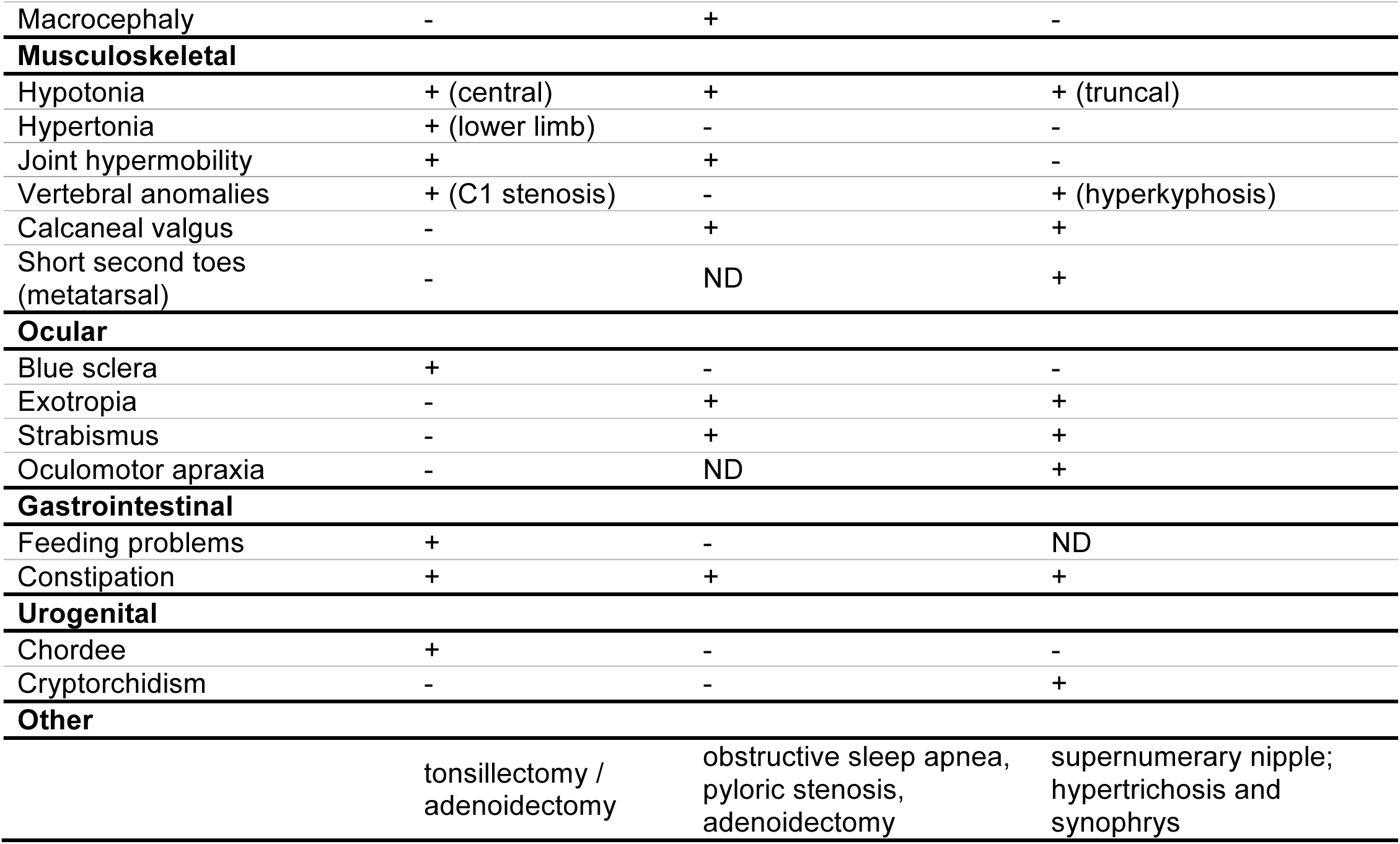
Mutations and Clinical Findings of Individuals with *KDM1A* mutations Plus (+) indicates presence of a finding, minus (−) indicates absence of a finding, C = central; LL = lower limb. ND = no data were available. N/A = not applicable. GERP = Genomic Evolutionary Rate Profiling. CADD = Combined Annotation Dependent Depletion. cDNA positions provided as named by the HGVS MutNomen web tool relative to NM_001009999.2

Variants with an alternate allele frequency >0.005 in EVS, 1000 Genomes, or ExAC, or >0.05 in an internal exome database of ~700 individuals were excluded prior to analysis. Additionally, variants that were flagged as low quality or potential false positives (quality score ≤ 30, long homopolymer run > 5, low quality by depth < 5, within a cluster of SNPs) were also excluded from analysis. Variants that were only flagged by the strand bias filter flag (strand bias > -0.10) were included in further analyses as the strand bias flag has previously been found to be applied to valid variants. Variants were annotated with the SeattleSeq138 Annotation Server, and variants for which the only functional prediction label was any one of “intergenic,” “coding-synonymous,” “utr,” “near-gene,” or “intron” were excluded. Individual genotypes with depth<4 or genotype quality<20 were treated as missing in analysis.

Code to generate Figure 3 is available at: http://dx.doi.org/10.6084/m9.figshare.1537555.

**Figure 3.** Effects of increasing number of trios sequenced and specificity of phenotype on power to detect significant association between putative mutations and phenotype. Assuming a de novo missense rate of 3.46×10^−5^/chromosome, as increasing numbers of trios (xaxis) are tested by exome sequencing, the power to detect a significant association (ranges of possible p values represented by different shades of green; darker indicates smaller and more significant p values) between de novo variants in a gene and the phenotype of interest increases. Additionally, as the specificity of the phenotype of interest increases, the proportion of individuals tested who have the phenotype (y-axis) naturally decreases, also resulting in increased power. A small decrease (60% to 50%) in the proportion of individuals who have the phenotype of interest can increase power more than sequencing 10,000 additional trios.

## RESULTS

Analysis of variants from ES under a *de novo* mutation model confirmed the presence of a different *de novo* variant in *KDM1A* (Refseq NM_001009999.2) in each of Families A and B (Table 1 and Figure 1; Figure 2). A complete phenotypic description, facial photographs, and published variant information from Family C were shared with the UW-CMG by the corresponding author (A.R.). All three children with *de novo KDM1A* variants had similar, albeit non-specific, clinical findings (Table 1) including similar facial features, global developmental delay and hypotonia (Figure 1; Table 1). In particular, all three individuals share a prominent forehead, slightly arched eyebrows, elongated palpebral fissures, a wide nasal bridge, thin lips, and wide-spaced teeth (Figure 1).

All three variants in *KDM1A* were missense variants predicted to be deleterious (minimum Polyphen-2 HumVar score of 0.962), result in amino acid substitutions of highly conserved amino acid residues (minimum GERP score was 5.72) in *KDM1A*, and have high CADD scores suggestive of dominant mutations (minimum CADD score 27.2) (Table 1). Moreover, *KDM1A* is in the top 2% of evolutionarily constrained genes, i.e. genes that are intolerant to functional variation, and this set of genes is enriched for genes known to underlie dominant Mendelian phenotypes^19^. None of the three variants were found in over 71,000 control exomes comprised of the ESP6500, 1000 Genomes phase 1 (Nov 2010 release), internal databases (>1,400 chromosomes), or ExAC (October 20, 2014 release). No rare variants in *KDM1A* were present in individuals included in Geno2MP v1.0^1^ who had a similar phenotype.

## DISCUSSION

### Function of KDM1A and delineating a new disorder

Tunovic et al.^16^ hypothesized that the phenotype of the proband in Family A might result from the combined effects of the *de novo* variant, c.2353T>C [p.(Tyr785His)], in *KDM1A* and a second *de novo* variant, c.2606_2608delAGA [p.(Lys869del)], in *ANKRD11*, i.e., suggesting that the child was affected by two Mendelian conditions, a Kabuki syndrome-like phenotype caused by the variant in *KDM1A,* and KBG syndrome (MIM 158050) caused by the variant in *ANKRD11*. This hypothesis was motivated, in part, by the presence of physical findings that did not overlap with features observed in Kabuki syndrome (MIM PS147920). However, comparison with two additional persons with *de novo* mutations in *KDM1A* reveals many of these features appear to be shared among all three. This suggests that mutations in *KDM1A* cause a condition that has phenotypic overlap with Kabuki syndrome but is nonetheless distinct.

Additional evidence suggests that the c.2606_2608delAGA [p.(Lys869del)] variant in *ANKRD11* in Family A does not cause KBG syndrome. Excluding microdeletions or large chromosomal deletions, the vast majority of *ANKRD11* variants that underlie KBG syndrome are frameshifts or nonsense mutations that are predicted to result in a truncated protein or nonsense-mediated decay^20-22^. In contrast, only four missense or small deletion/duplication mutations, have been reported as causing KBG syndrome^20-22^. These findings, combined with the observation that *ANKRD11* is not highly conserved (only 79% identity with its mouse ortholog^21^) and is highly polymorphic in the general population^23^), suggest that only a small subset of missense variants found in *ANKRD11* result in KBG syndrome. Moreover the c.2606_2608delAGA [p.(Lys869del)] variant is not predicted to be pathogenic by CADD v1.0 (a Phred-scaled score of 13.03 is well below the score of 25 observed for the majority of mutations that cause autosomal dominant conditions). Finally, although macrodontia of the upper incisors, considered a hallmark feature of KBG syndrome, is often not observed until adult teeth emerge, the proband in Family A has normal dentition^20,22,24^.

KDM1A is a histone demethylase that has been extensively studied *in vitro* and in model organisms and has been shown to play diverse and key roles in regulating gene expression during development.^25^ Homozygous knockout of *Kdm1a* in mice is lethal during early embryogenesis^26^. Kdm1a is involved in repression of neuronal genes in non-neuronal cells^27,28^, and during the perinatal period, alternative splicing of KDM1A results in expression of two neuron-specific isoforms that regulate neurite maturation^29^. In mice, proper skeletal muscle differentiation requires Kdm1a to demethylate myogenic promoters^30^, which may explain the discovery of heart defects in mice homozygous for a hypomorphic *Kdm1a* allele^31^. Interestingly, mice heterozygous for a *Kdm1a* deletion are apparently normal and fertile^26^, suggesting that haploinsufficiency may not result in an obvious defect. Nevertheless, it remains to be seen if the phenotypes we report to be caused by variants in *KDM1A* are due to loss or gain of function. An additional intriguing observation is that all three mutations alter residues in the amine-oxidase domain (Figure 2B), which is composed of FAD-binding and substrate-binding functional subdomains.^32^ The active site cavity of KDM1A is formed by the substrate-binding subdomain^32^ and is required for KDM1A to demethylate H3K4me1/2 and repress transcription^28^.

### Scaling up gene discovery by social networking to tackle the n-of-1 problem

The discovery that variants in *KDM1A* underlie a distinctive and previously unrecognized Mendelian condition is the result of social networking by the family of an affected child with another family and several researcher groups. This approach consisted of establishing a website that included a comprehensive description of the proband’s symptoms and medical history, using both lay and medical terminology and reports of putative pathogenic variants identified via ES, and a linked blog, Twitter account, and Facebook page. Exposure to the public-at-large via common social media such as Facebook and Twitter is a strategy that leverages sites familiar to many families. Technology-savvy families are also capitalizing on existing searchable information platforms such as editing entries for conditions described on Wikipedia, setting Google alerts for symptoms and rare variants, purchasing Google adwords, and using Google analytics to identify pockets of researcher and patient activity.^12^ In this case, only five days after launching their website, Family A received an email from Family B describing “her son, along with a picture of him that showed the remarkable resemblance between the two boys – he looked like he could be [proband A]’s brother” (P. Lorentzen, personal communication).

The direct-to-consumer genetic testing movement, particularly genetic ancestry testing, has made it routine to use of the internet and social media to research genetic relationships and the meaning of genetic information, including variants associated with disease.^33^ Building on the global reach of the internet, online social networking is also increasingly leveraged by rare disease communities to connect families, enabling them to share their experiences, provider/researcher relationships, genetic knowledge, and strategies for advocating for their children.^34-36^ Indeed, one of the important benefits of social networking is the ability of families to share information, including sequence data, directly with researchers in hopes of garnering more interest and making collaboration more convenient and cost-effective. Accordingly, online social networking is increasing the role that families play in stimulating, coordinating, and supporting research.

To support and facilitate the efforts of families toward discovering the genetic basis of their condition, we developed an online portal, the Repository for Mendelian Disorders Family Portal (RMD-FP) for families to submit and subsequently share phenotypic information and ES/WGS data. The RMD-FP is a point of entry into the human genetics community for families who seek to cast the widest net in recruiting researchers to work on their condition. The RMDFP provides information to families about research, facilitates family decisions about their preferences for how their data may be used, and guides families through the process of directly submitting their phenotypic information and genomic data. The RMD-FP will eventually enable collection of detailed self-reported phenotype/trait information via structured data entry and enable families to receive results, if available, via My46, a web-based tool for managing return of genetic test results.

Once data are deposited in the RMD-FP, the phenotypic information will be curated and structured^37^ for submission to PhenomeCentral/Matchmaker Exchange. Genomic data will be re-analyzed and all variants found by either prior diagnostic sequencing or re-analysis to be segregating under the appropriate inheritance model(s) (i.e., candidate variants) will be entered along with the structured phenotypic data into a database. If a small number of candidate genes are identified, the genes will also be submitted to GeneMatcher/Matchmaker Exchange. However, many families consist of a single affected individual with no contributing family history, leading to analysis under all possible standard inheritance models (e.g., homozygous recessive, compound heterozygous, and *de novo*). This results in a large list of candidate variants/genes that are not appropriate for submission to GeneMatcher, and currently leaves families no way to efficiently share candidate variants with other interested families, clinicians, diagnostic laboratories, and researchers. The combination of structured phenotype information and sharing of all candidate variants should increase data consistency and thus the probability of a match.

To help address this gap, we are developing MyGene2 (beta release projected early 2016), a public web-based tool that enables searches of candidate variants/genes linked to phenotypic profiles of persons and families deposited in the RMD-FP. Users of MyGene2 will be able to search for candidate variants matching a gene, inheritance model, and/or phenotypic trait or profile. If a user identifies a variant of interest, they can register with the site by creating an account, which enables them to anonymously contact the submitter(s) for further information. Registration is required to protect the confidentiality of sample submitters, to track matches, and to survey matched users about subsequent discoveries and publication. Tracking outcomes also helps to ensure that families benefit from their participation. Families will be able to use sample submission to publicize their candidate variants/genes, make their data available to the community via an editable “family page,” and participate more fully in gene discovery efforts without requiring a high level of technical knowledge. Clinicians and researchers will also be able to search for additional families with mutations in the same gene for gene discovery and delineation of new conditions; and diagnostic laboratories will be able to search for additional cases to assist in interpretation of VUS. Matches made through MyGene2 will only be required to acknowledge the site and its sources of support in ensuing manuscripts. Indeed, we envision MyGene2 as a resource to empower families, clinicians and investigators to delineate new Mendelian conditions largely independent of UW-CMG so as to accelerate the rate of gene discovery for Mendelian conditions. With broad participation from the human genetics community, MyGene2 has the potential to greatly facilitate overcoming the “n-of-1 problem”.

The scenario we report in which a variants of unknown significance in the same candidate gene led to ascertainment of several persons with overlapping clinical features and delineation of a distinct syndrome is likely to become an increasingly common strategy of discovering genes for Mendelian conditions. Identification of three independent families in which each person with a de novo variant in the same gene has the same condition meets existing guidelines for causality of Mendelian disorders.^1,3^ Nevertheless, confidence would be gained by assigning a p value for this observation^38^, but doing so is difficult in the absence of greater sharing of detailed phenotype information linked to ES data from a large number of independent cases. For example, developmental delay is perhaps the most common phenotype found in individuals that undergo clinical ES, comprising 64% of cases tested in one recent survey.^39^ Therefore, if we estimate that roughly 10,000 trios have been analyzed for *de novo* variants via clinical ES, ~6,400 are predicted to have had developmental delay^39^. Using a Fisher’s exact test for independence between the presence of de novo variants in *KDM1A* and developmental delay, the p value for identifying three individuals with *de novo* variants in *KDM1A* and developmental delay and no individuals with de novo variants in *KDM1A* without developmental delay is only 0.5576. This p-value is not significant because of poor statistical power to distinguish between persons with mutations in different genes who are broadly described as having the same common, non-specific condition.

Power can be improved by increasing the sample size of trios tested; by using additional phenotypic details to increase specificity about the phenotype tested; or some combination thereof, with increasing specificity of the phenotype of interest being much more efficient at improving power (Figure 3). For example, if we consider a gene with a de novo mutation rate of 3.46×10^−5^ per chromosome (i.e. the mean predicted de novo mutation rate for missense mutations in the top ~5% of highly evolutionarily constrained genes in a recent study^19^), even if 100,000 trios are tested, the best p value that could be obtained as long as 64% have developmental delay is 0.094. In contrast, if we increase the specificity of the phenotype of interest and thus reduce the fraction of those same 100,000 cases with the phenotype to 50%, the p value drops to 0.016. While increasing both sample size and phenotypic specificity is ideal, a quick and effective way maximize power of existing datasets is to make deep, structured phenotypic data linked to genotype data publicly available and accessible via tools like MyGene2 and others in order to enable statistically-rigorous assessment of similar Mendelian gene discoveries.^3,38^

Mendelian gene discovery has traditionally been organized around the clinician-researcher as the central hub, around which families are solicited, experiments performed, results reported in manuscripts, and data shared. Social networking promotes a more egalitarian network in which families also act as nodes, independently sharing phenotypic information, genetic data, and results with other families and researchers.^14^ Yet, this can be a labor-intensive, inefficient, and expensive endeavor that requires some technical expertise to maximize the effort. At its full potential, MyGene2 can serve as an organizing node for families, providing them with convenient and free access to data from a large number of other families and investigators. It should be noted that MyGene2 is but one new tool to facilitate social networking and data sharing among persons interested in rare diseases. We expect and indeed encourage development of other strategies and solutions^40^ toward the same goals. One outcome of this model is a somewhat diminished role for both clinicians and investigators as the central organizing node but greater empowerment of families, and we predict, a greater rate of discovery of genes and newly-delineated Mendelian conditions.

In summary, social networking among families led to the recognition, if not the frank discovery, that de novo variants in *KDM1A* cause a newly-delineated condition characterized by developmental delay, hypotonia, and characteristic facial features. Coupled with the fairly narrow range of phenotypic variation observed in the affected individuals described herein, it is likely that mutations in *KDM1A* might also explain some cases of apparently isolated intellectual disability. Developing infrastructure to empower families to share phenotypic information and genetic data at scale would empower many more families worldwide and we predict will accelerate the pace of gene discovery for Mendelian conditions. The rapid translation of these discoveries into diagnostic tests and new starting points for repurposing or developing therapeutics would, in turn, improve the overall care of families with rare diseases.

## SUPPLEMENTARY MATERIAL

Supplemental Data includes 1 figure.

## DISCLOSURE

M.J.B., H.K.T., and J-H.Y. have a patent application pending on My46. The other authors declare no conflict of interest.

## ACKNOWLEDGEMENTS

We thank the families for their participation and support and James Barkovich, Anne M. Slavotinek, Elliott H. Sherr, and Brent M. Werness for helpful discussion. Our work was supported in part by grants from the National Institutes of Health / National Human Genome Research Institute and the National Heart, Lung and Blood Institute (1U54HG006493 to M.B., D.N.; 1RC2HG005608 to M.B., D.N.; 5R000HG004316 to H.K.T.), National Institute of Child Health and Human Development (1R01HD048895 to M.J.B.), the Life Sciences Discovery Fund (2065508 and 0905001), and the Washington Research Foundation. The authors would like to thank the Exome Aggregation Consortium and the groups that provided exome variant data for comparison. A full list of contributing groups can be found at http://exac.broadinstitute.org/about. The authors would like to thank the University of Washington Center for Mendelian Genomics and all contributors to Geno2MP for use of data included in Geno2MP.

## Web resources

The URLs for data presented herein are as follows:

MyGene2: http://www.mygene2.org

Repository for Mendelian Genomics Family Portal: http://uwcmg.org/#/family

Exome Variant Server (NHLBI Exome Sequencing Project ESP6500): http://evs.gs.washington.edu/EVS/

Exome Aggregation Consortium (ExAC), Cambridge, MA [accessed October 2014]: http://exac.broadinstitute.org

Geno_2_MP, NHGRI/NHLBI University of Washington-Center for Mendelian Genomics (UWCMG), Seattle, WA [accessed January 2015]: http://geno2mp.gs.washington.edu

GenomeConnect: http://genomeconnect.org

GeneMatcher: http://genematcher.org

Human Genome Variation: http://www.hgvs.org/mutnomen/

Matchmaker Exchange: http://matchmakerexchange.org

Milo’s Journey: http://milosjourney.com

PhenomeCentral: http://phenomecentral.org

Online Mendelian Inheritance in Man (OMIM): http://www.omim.org/

SeattleSeq: http://snp.gs.washington.edu/

